# Non-targeted Screening of Commercial CBD Products in the United States Reveals Common Contamination and Adulteration

**DOI:** 10.1101/2020.08.25.267302

**Authors:** Ben Orsburn

## Abstract

The production of hemp and products derived from these plants that contain zero to trace amounts of the psychoactive cannabinoid tetrahydrocannabidiol (THC) is a rapidly growing new market in the United States. The most common products today contain relatively high concentrations of the compound cannabidiol (CBD). Recent studies have investigated commercial CBD products using targeted assays and have found varying degrees of misrepresentation and contamination of these products. To expand on previous studies, we demonstrate the application of non-targeted screening by high resolution accurate mass spectrometry to more comprehensively identify potential adulterants and contaminants. We find evidence to support previous conclusions that CBD products are commonly misrepresented in terms of cannabinoid concentrations present. Specifically, we observe a wide variation in relative THC concentrations across the product tested, with some products containing 10-fold more relative signal than others. In addition, we find that several products appear to be purposely adulterated with over the counter drugs such as caffeine and melatonin. We also observe multiple small molecule contaminants that are typically linked to improper production or packaging methods in food or pharmaceutical production. Finally, we present high resolution accurate mass spectrometry data and tandem MS/MS fragments supporting the presence of trace amounts of fluorofentanyl in a single mail order CBD product. We conclude that the CBD industry would benefit from more robust testing regulations and that the cannabis testing industry, in general, would benefit from the use of non-targeted screening technologies.

## 1. Introduction

New products containing cannabidiol derived from cannabis plant material have appeared on shelves at a rapid pace in the United States following the decriminalization of hemp production in 2019.[1] These products are derived from cannabis plant materials in a variety of ways, typically through solvent or supercritical CO_2_ extraction. [*2–4*]. The central theme of these products is that they contain high concentrations of non-psychoactive cannabinoids, primarily cannabidiol (CBD). Again, while laws vary, all states currently require zero to negligible amounts of the psychoactive cannabinoid, tetrahydrocannabidiol (THC) and cannabinoids that can be instantly converted to psychoactive forms through processes as simple as heating the product.

While these products are relatively new to the U.S., they have been legal or decriminalized in some locations for much longer. A study of CBD oils commercially available in Europe by Pavlovic *et al*., in 2018 thoroughly analyzed 14 products using a variety of techniques, including high resolution accurate mass spectrometry (HRAM). This study found a high number of discrepancies in this small sample set, including inaccurate reports of cannabinoid concentrations in 9 of 14 products, and chemical evidence that did not support how the labels stated the products were produced.[5]

A study in the U.S. that focused exclusively on the concentrations of cannabinoids in online CBD products found of 84 products tested, only 30.4% were accurately labeled within the tolerances defined by the study.[6]

As an expansion on these earlier works, we applied non-targeted screening techniques by ultra-high pressure liquid chromatography coupled HRAM mass spectrometry (LCMS) to 21 commercially available CBD products. Non-targeted screening relies on the acquisition of chromatographic features and mass spectrometry data to examine samples in an unbiased way. Post-acquisition, algorithms are used to attempt to identify and apply relative quantification data to the ions that are seen.[7] Historically, non-targeted screening was only possible when spectral libraries of pure known compounds had been painstakingly created and manually curated. In these algorithms, software is used to simply look for high homology between the data acquired and the spectra in the libraries. More recently, new tools have shown the promise of identifying molecules that are not present in libraries by applying hybrid search techniques that rely on libraries but can adapt to mass shifts due to currently unknown chemical modifications.[8]

Unlike targeted mass spectrometry methods, non-targeted screening casts a wider net, allowing compounds to be identified in products with no *a priori* knowledge of their presence.[9,10]. While a wide range of potential contaminants may be rapidly identified with these techniques, validating the presence of compounds may be labor intensive. Informatics to automate and improve small molecule identification is a rapidly evolving field, driven largely the emerging field of metabolomics, which seeks to identify and characterize changes in the complete population of small molecule metabolites in biological systems.[11–13] Metabolomic studies may attempt to make matches from biological materials to spectral libraries containing thousands of potential compounds. One example of recent advances is the mzCloud online library. The mzCloud is a database of small molecule tandem mass spectra comprised entirely of HRAM tandem mass spectra acquired at multiple fragmentation energies. The high mass accuracy of this data allows more certainty in identifications by reducing the number of potential matches. It is important to note, however, that these techniques have well known limitations and extensive validation may be required to fully elucidate the identity of a small molecule. Although methods for statistically scoring false discovery rates have been developed they have not been fully implemented in routine non-targeted workflows.[14–16].

In this study we apply non-targeted screening to 21 commercially available CBD products. As in previous studies we find that cannabinoid content and other chemical evidence does not appear to be reflective of the product labels. Furthermore, we conclude that product adulteration appears to be widespread in the industry, with multiple products containing undocumented compounds, including over the counter and prescription drugs.

## 2. Results

### 2.1. Deliberately mislabled products

Although regulations in the U.S. have become more lenient regarding the use of hemp and CBD products, the Food and Drug Administration has issued nearly 20 warning letters to producers in this industry since 2019.[17,18] Two separate products purchased for this study from online retailers indicated on their labels that they were approved for use by the FDA, statements that are not true. Examples of these labels are shown in Figure 1.

**Figure 1.**
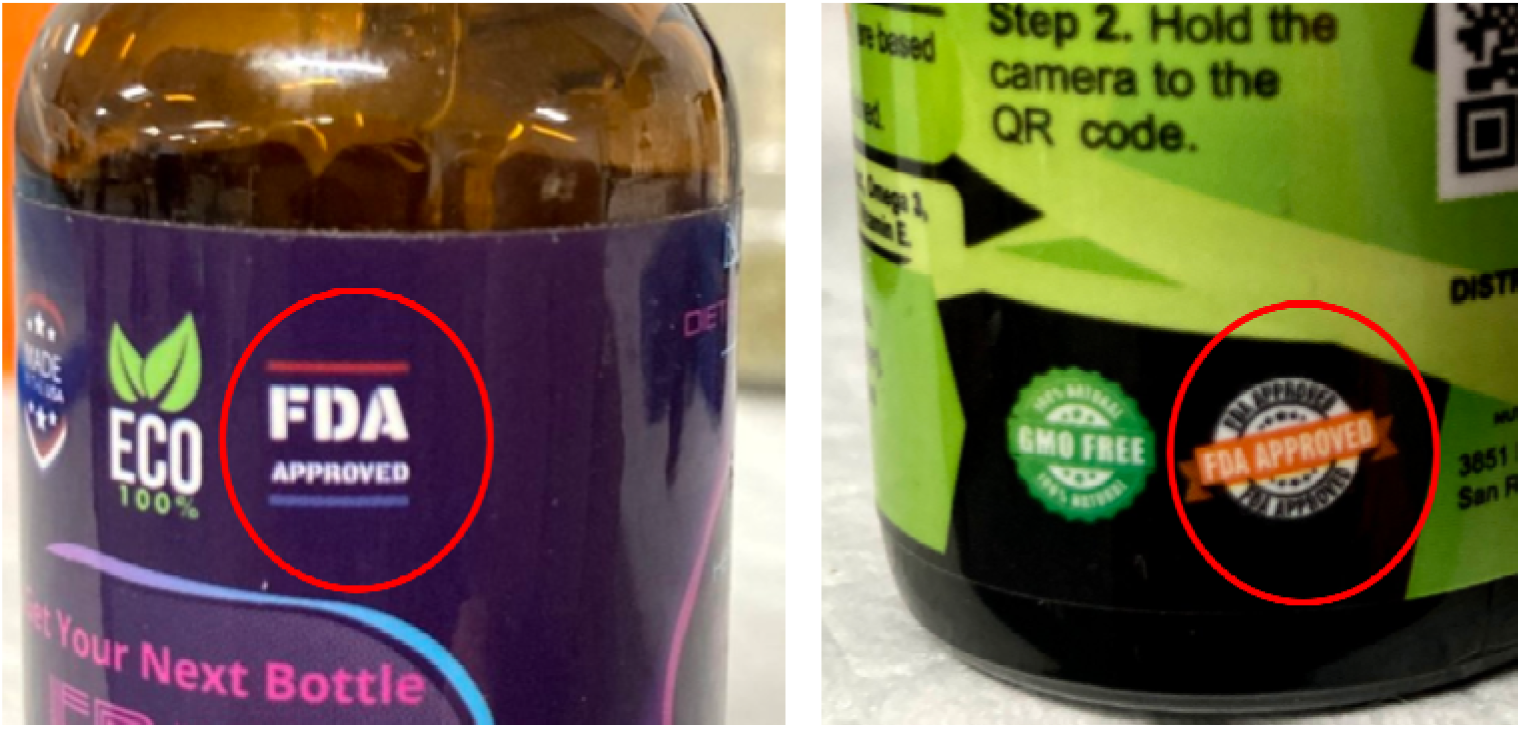
Two example products with misrepresentations stating the endorsement of the Food and Drug Administration on their respective contents.

### 2.2. Cannabinoid Relative Concentrations

The design of this study was exploratory in nature and absolute quantification of cannabinoids is outside of the scope of the work presented here. From the relative quantification data we can see a high degree of fluctuation in cannabinoid content across products. Figure 2 is the extracted ion chromatogram from products that were found to possess THC above our limit of detection. By comparing the areas under the curve of these product signals we can obtain a relative metric of the THC concentration. While most products in our study did not have quantifiable levels of THC, we find a 6-fold variation in relative THC content in the ones that do, likely pushing some products over the federally mandated maximum of 0.03%.[19]

**Figure 2.**
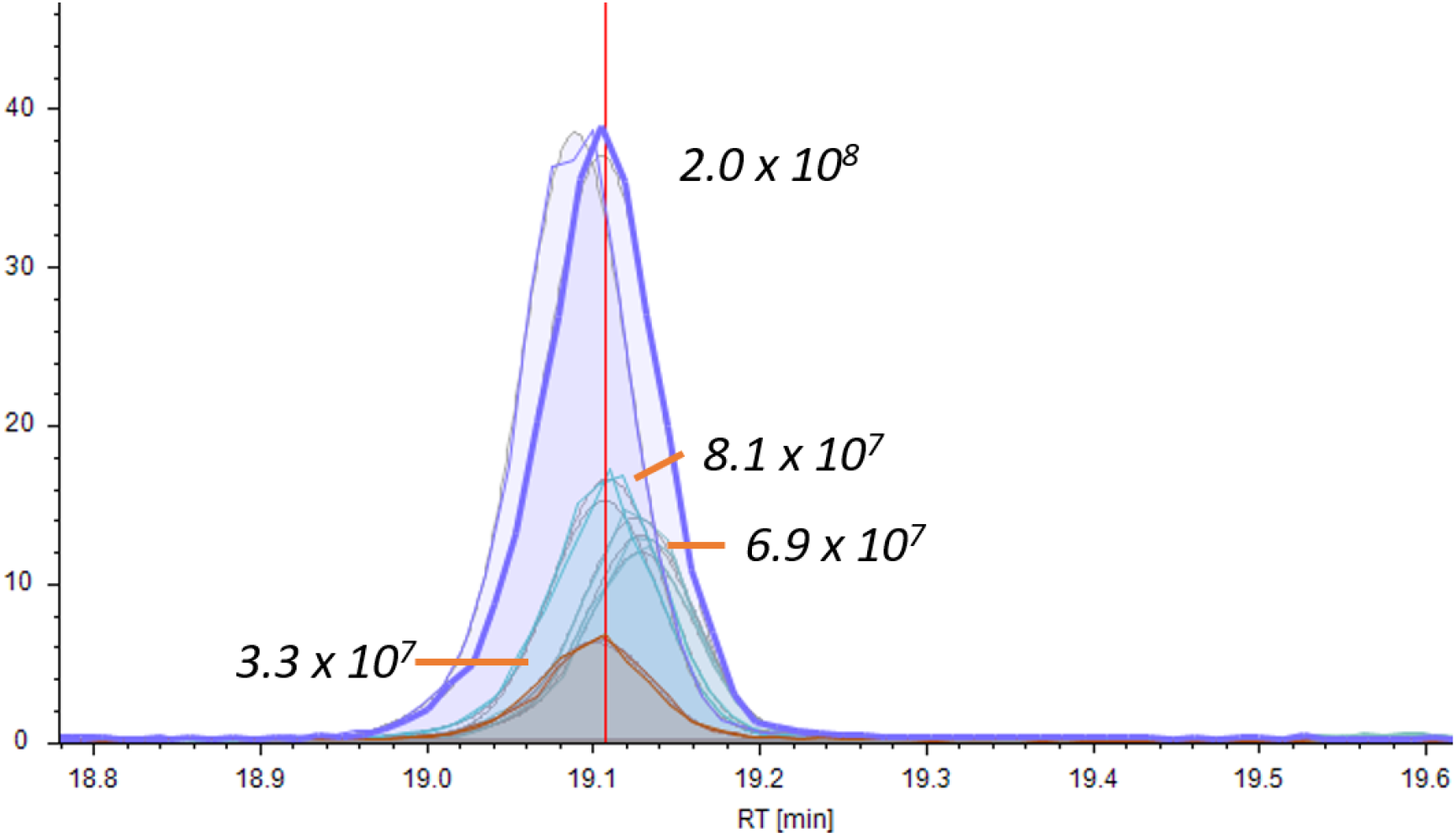
Visualization of the LCMS peak for THC in 4 products above the limit of detection (LOD) for the molecule. The relative area of each peak is presented as the results of 3 technical replicates.

### 2.3. Identification of Adulterants

Acquired data was searched against the libraries summarized in Table 1. A common method in non-targeted screening is the utilization of multiple databases. Confidence can be established based on obtaining multiple matching identifications from complementary libraries. Furthermore, no library today is comprehensive and drawing from multiple sources increases the likelihood of identifying a compound.[20,21]

**Table 1.** A summary of the databases employed with version identifier, if applicable, or date library downloaded or accessed for processing these files.

| Library | Approximate number of compounds |
| --- | --- |
| Drug Bank 5.1.6 | 7,800 |
| Cayman Chemical (8/10/20) | 16,244 |
| Arita Flavonoid Library [22] | 6,300 |
| Extractables and Leachables Library [23] | 1,544 |
| Environmental Food Safety Library [19] | 1,620 |
| mzCloud (8/10/20) [24] | 18,200 |

Due to the high mass accuracy of both the MS1 and MS/MS measurements present in the mzCloud database, we choose to primarily rely on high confidence hits obtained from this source. Compounds are readily verified by manually comparing the experimentally obtained data to spectra from these libraries. Figure 3 is an example of one such output for the identified prescription medication Valpromide.

**Figure 3.**
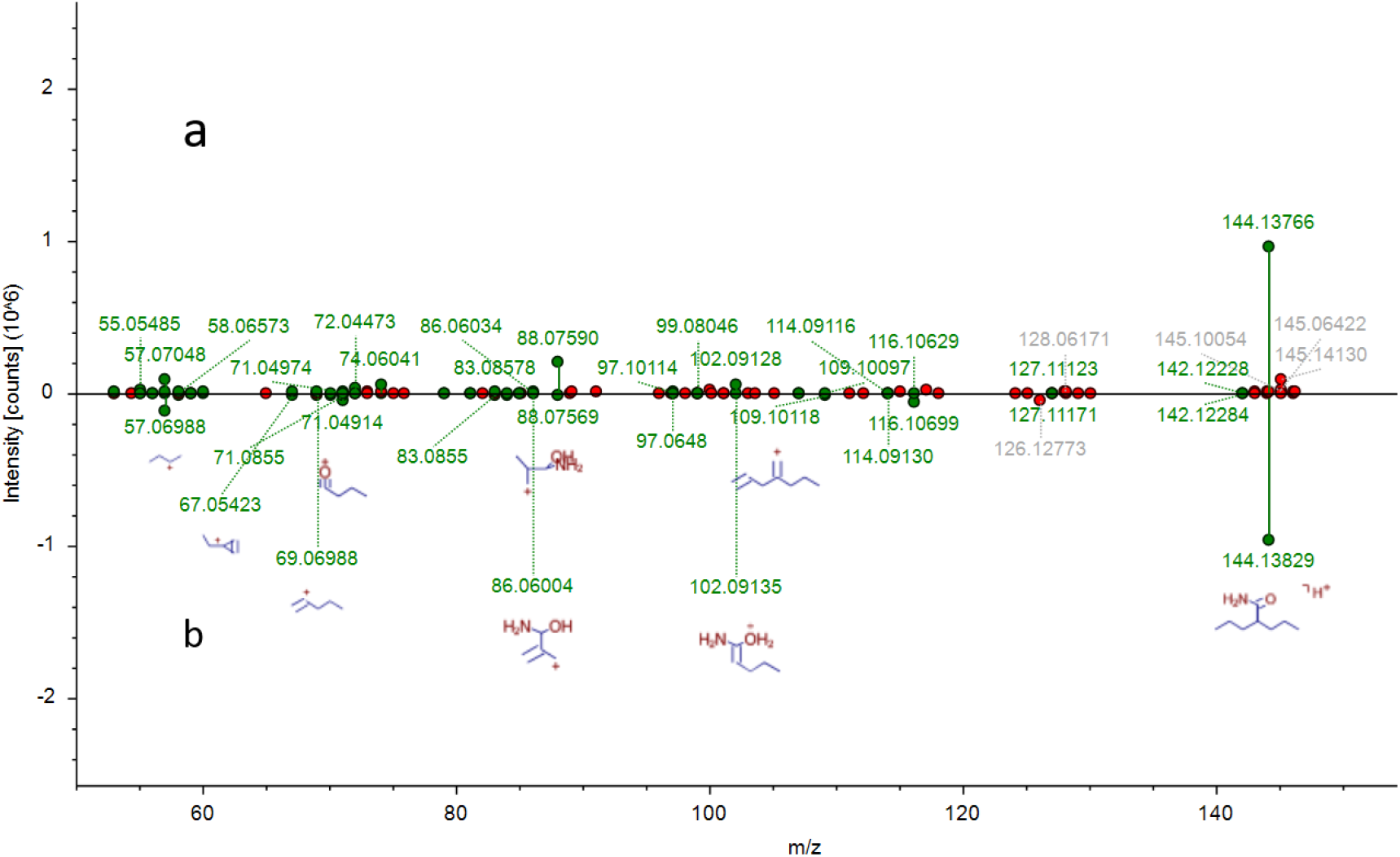
Example mzCloud match data for a compound flagged as having 91.0% match for Valpromide. (a) The experimentally obtained data from the mail-order CBD product analyzed (b) The mzCloud MS/MS spectra. Fragment ions that match are flagged in green and identified product ion structures are shown if available in the library. Fragments observed in the experiment but not in the library are labeled in red.

The LCMS small molecule profile of any material is complex, and many ions are often observed that elute chromatographically with similar mass to charge ratios (m/z). A common practice in mass spectrometry is the use of smaller isolation windows [26] and multistage fragmentation to reduce interference from coeluting molecules.[27] While these approaches help, co-fragmentation is a feature of mass spectrometry that has no current solution. We can see in Figure 3 that several lower intensity ions higher in mass than the targeted parent were coisolated in this spectrum and they undoubtedly contribute to the unidentified fragment ions displayed. However, due to the complete justification of the fragments we can conclude that that this compound identification is one that should be flagged for targeted validation. An increasing number of matching fragment identified increases confidence in identifications as the likelihood of these fragments occurring at random, or in other compounds correspondingly decreases.[10,28–31] A second example of an mzCloud match is shown Figure 4. The spectra shown was identified by mzCloud and Compound Discoverer as a match for 3-Fluorofentanyl. Figure 4 has been rescaled to better visualize the number of ion fragments that support the strength of this match.

**Figure 4.**
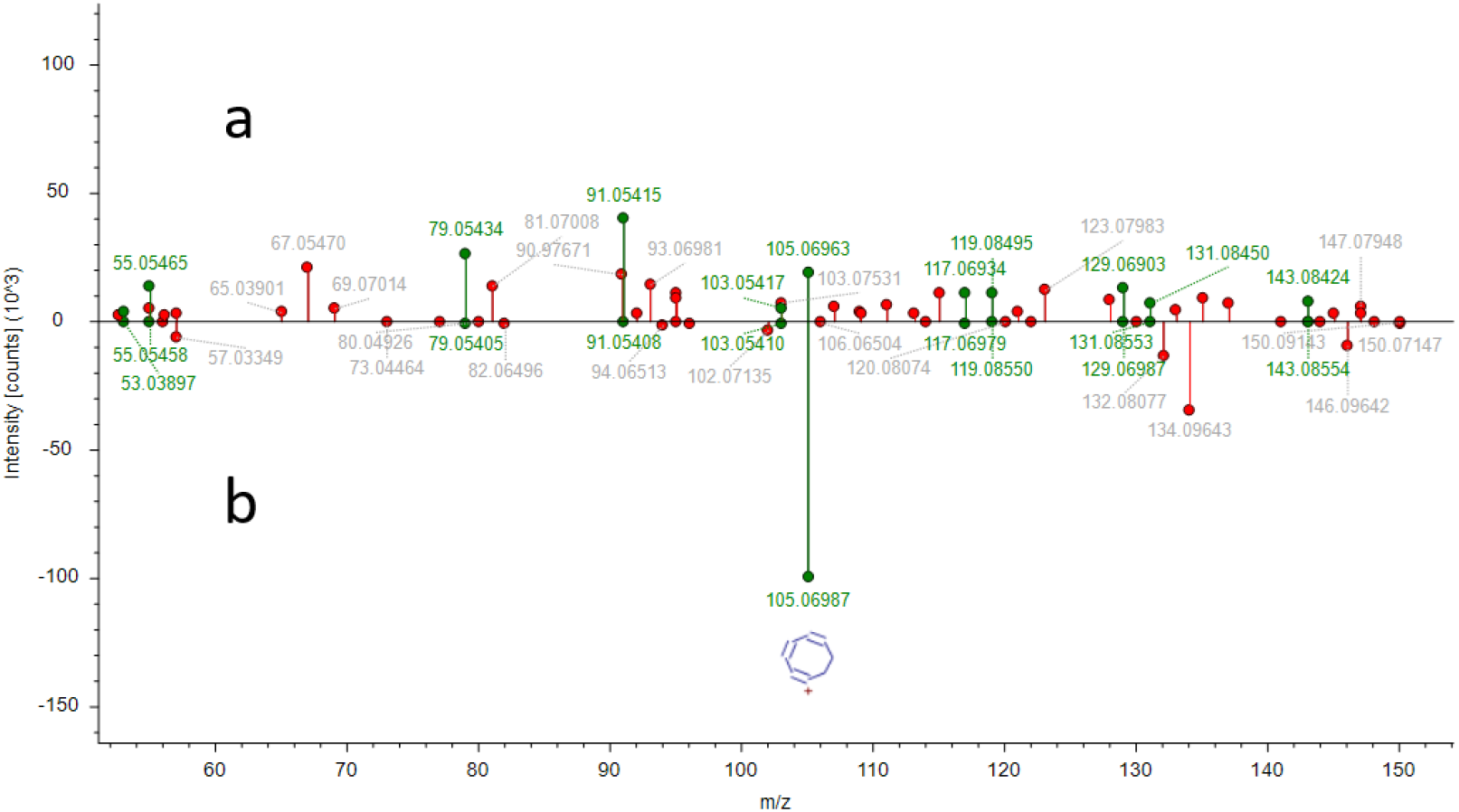
Mirror plot displaying the mzCloud predicted match to 3-Fluorofentanyl for MS/MS spectra zoomed to display the highest density of matching fragment ions. (a) experimental MS/MS spectrum (b) mzCloud library spectrum.

A recent LCMS analysis of fentanyl compounds carried out Strayer *et al*., optimized methods for the identification of 24 illicitly manufactured fentanyl (IMF) analogues.[32] The resulting method relies on finely optimized chromatographic conditions and fragmentation energy as fentanyl analogues have highly similar retention characteristics on reversed phase chromatography and therefore commonly coelute. IMF available on the black market is often a blend of multiple analogues due to crude illicit manufacturing practices.[33–36]. Although IMF analogues are a diverse class of compounds, they possess a shared basic core structure. In the Strayer *et al*., study quantitative and qualifier ions were optimized for identifying 24 fentanyl compounds in complex matrices. Of the 48 fragment ions chosen, 14 are the core diagnostic fragment shown in Figure 4, corresponding to ~105.070 m/z. Fentanyl analogues have been thoroughly studied with HRAM mass spectrometry and the release of a diagnostic ion with a corresponding *m/z* to this ion is used to help confirm the presence of IMF in commercial products and patient samples, demonstrating the specificity of a high mass accuracy spectrum displaying this ion.[37–39]

### 2.3. Product summaries and compound identifications

A summary of other compounds identified with high confidence is presented in Table 2. Of the 21 products studied in this phase, eleven appear to be affected by adulteration, contamination or misrepresentation. The most common contaminant identified, Erucamide, is a surfactant used in manufacturing and is a well-charactarized extractable/leachable contaminant in pharmaceutical products.[23] Polyethylene glycol n14 and n15 were identified with high confidence in two separate materials, as was the over the counter sleep additive melatonin. Supplemental Information I provides additional support for these identifications by providing the mirror plot for each compound in Table 2.

**Table 2.** Compounds identified in some of the products tested.

| Compound | Found in samples | Classifications |
| --- | --- | --- |
| Caffeine | FS9 | Adulterant |
| Valpromide | FS1 | Adulterant |
| Melatonin | FS10,FS7 | Adulterant |
| Theobromine | FS9 | Adulterant |
| 1-Stearoylglycerol | G1 | Contaminant |
| Polyethylene Glycol N14 | B4,B5 | Contaminant/Extractable |
| Polyethylene Glycol N15 | B4,B5 | Contaminant/Extractable |
| Polyethylene Glycol N10 | G1 | Contaminant/Extractable |
| 1-Naphthol | FS11 | Contaminant/Extractable |
| Nootkatone | G1 | Contaminant |
| Palmitoleic acid | G1 | Adulterant |
| Yohimbine | FS9 | Adulterant |
| Elevated relative THC | B4 | Misclassification |
| Erucamide | K12,K270,FS5,FS7, | Contaminant |
| Tributyl Phosphate | B4 | Extractable/Leachable |
| Proposed Fluorofentanyl | B4 | Adulterant/Drug of Abuse |

## 3. Discussion

In this study we have utilized well-characterized techniques for non-targeted screening to evaluate 21 commercially available CBD products. This is a logical extension on previous studies that have evaluated similar products both in the U.S.[6] and in Europe.[5] What we have found in the application of non-targeted screening is similar in many regards to the results of the studies this work is based on. There are clearly good actors in this emerging market, as 10 of 21 products we have studied revealed no obvious contaminations or surprise compounds. However, simply observing the labeling and advertising of some commercially available products is enough to conclude that there are also bad actors in this industry. Using a standard non-targeted screening assay in just this small sample set we have identified 6 products that may have been purposely adulterated with compounds of known activity in humans. While caffeine and melatonin are commonly used and typically harmless, accurate product labeling is essential for the protection of members of the population that are not tolerant of specific product ingredients. More nefarious is the potential identification of a fentanyl analogue, as well as the potential identification of the prescription medication Valpromide. Fentanyl is potent, highly addictive and was the likely cause of 30,000 deaths by overdose in the US in 2018.[34] Valpromide is a medication used for psychiatric disorders, the prevention of epileptic seizures and has been characterized as increasing the activity of other drug classes.[40] As noted in a previous study [6] the potency of CBD products is often under reported. One product in this study that demonstrates the highest levels of contamination and adulteration, also appears to possess elevated levels of THC compared to our control samples.

It should be noted that non-targeted screening techniques make identifications that should be carefully examined with validated techniques.[7] To conclusively verify the identity of these compounds, despite the level of confidence match to libraries, we must obtain pure reference standard materials to prove that the compounds of interest match in elution profile and fragmentation on our specific systems.[41] Furthermore, proving the presence of 3-Fluorofentanyl in the sample described would require the cooperation of a DEA certified lab with clearance to purchase, possess and use this material as a control standard.[42] Validating the identifications presented herein are therefore beyond the scope of this current study.

We do, however, see reason for concern in this new marketplace and believe that non-targeted screening by HRAM mass spectrometry could play a valuable role in protecting consumers. While cannabis products with higher levels of psychoactive cannabinoids are regulated in many states for potency and potential contaminants, CBD products are not being evaluated with such scrutiny. Furthermore, in an informal survey of professionals in the cannabis testing industry we were only able to identify one lab in the U.S. that uses HRAM mass spectrometry and therefore has the capacity for non-targeted screening as presented here (informal results not presented here). Initiating non-targeted screening as a tool in the emerging cannabis industry may prove to a be challenge, but one we must conclude would be worth the rewards in protecting consumers.

## 4. Materials and Methods

### Preparation of samples for LCMS

Approximately 0.25 grams of each sample transferred into a 15 mL graduated centrifuge tube (Azer Scientific) and the actual mass was recorded for later normalization. Each sample was diluted with 10 mL of LCMS grade methanol with the final mass of solvent likewise measured by mass. The mixture was briefly vortexed to ensure basic homogenization and placed in a sonic water bath and sonicated for 15 minutes at room temperature. The materials were allowed 20 minutes to settle at room temperature. Approximately 20 μL of the supernatant was transferred to a gastight screwcap autosampler tube (Fisher Scientific) for untargeted GCMS analysis (data not presented here). Approximately 1 mL was removed by sterile disposable syringe and filtered through a 0.22 μm nylon syringe filter directly into a 2 mL amber glass autosampler vial (Azer Scientific) for LCMS analysis

### LC-MS Analysis

All LCMS analysis was performed on Vanquish uHPLC (Dionex) coupled to a Q Exactive mass spectrometer (Thermo Fisher). A 2 μL injection of each sample was utilized. Samples were crudely cleaned prior to injection onto the analytical column using a proprietary 30 second solid phase extraction process inline (CRL) prior to moving the sample to 10 cm C-18 HPLC column. For discovery runs the reversed phase gradient ramped from 2-5% buffer A (100mM ammonium formate, 0.1% formic acid in LCMS grade water) to 60% buffer B (100mM ammonium formate, 0.1% formic acid in LCMS grade methanol) in 7 minutes followed by an increase to 95% B by 15 minutes. These conditions were maintained for 3 minutes before returning to baseline conditions. Throughout the experiment, a flow rate of 0.3 mL/min was maintained. For quantitative experiments not presented here, all LC conditions were identical, with the exception that the baseline condition was 0% buffer A to facilitate the effective solid phase extraction of plant growth regulators to meet PA Title 28 1171 regulations. The Q Exactive operated exclusively in positive ionization mode using the following source conditions: ESI Voltage 3500V, Capillary Temperature 275, Sheath gas 40 (arbitrary units), Aux Gas 10 (arbitrary units), Spare Gas 1.5 (arbitrary units), Probe heater temperature 250C, S-Lens RF 50.0%. The Q Exactive was operated in data dependent mode with an MS1 from 110-1000 m/z at 70,000 resolution, with an AGC target of 1e6 charges and a maximum injection time of 200ms. The 3 most intense ions were selected for fragmentation using a 4.0 Da isolation window. The isolated ions were fragmented using stepped collision energy normalized to a +1 default charge and NCE of 20, 35, and 100. The MS/MS spectra are acquired from the acquired fragmentation at all 3 energies simultaneously. The MS/MS spectra were recorded at a resolution of 17,500 at a reference mass of 200 m/z.

### Data Processing

All vendor .RAW files were processed in Compound Discoverer 3.1.0.305. Briefly, all technical duplicates were grouped together under a single identifier. All full MS1 and MS/MS scans were identified by the Select Spectra Node using MS(n-1) precursor selection. Retention times were aligned using an Adaptive Curve model with a maximum 3 minute shift and 5ppm mass accuracy cutoff. Adducts were automatically detected and condensed using all 33 adducts provided by the detect compound nodes. Adducts that are not possible under the sample preparation and analysis conditions are utilized to help establish a metric for the number of false discoveries identified in the analysis. Compounds were grouped at 5ppm mass tolerance and a 0.2 minute retention time tolerance post alignment, with the preferred ion adducts [M+H]+1. The grouped compounds were analyzed in parallel with 3 different programs before the results were combined. 1) Molecular compositions were predicted using the vendor default parameters of 5ppm mass accuracy, and only using isotopes with a signal to noise threshold greater than 3-fold. When available fragmentation das utilized for candidate ranking with the same tolerances described for MS1. 2) All compounds were searched against in-house developed database containing a library of known pesticides, plant flavonoids and common chemical contaminants in food processing using an MS1 mass tolerance of 5ppm and a retention time tolerance of 2 minutes where applicable. 3) All compounds with MS/MS data were searched against the mzCloud database using libraries using 17 databases for drugs and chemical contaminants. To improve the confidence of mzCloud matches, the collision energy was matched with tolerance of 20ev based on the center collision energy in the study of 58eV NCE with a fragment intensity threshold applied. Precursor and fragment tolerance for all scans was set at 10ppm.

## Acknowledgments

The author would like to thank Meghan Pettis, Kelsey Schubert, Stanley Keyton and David Berryhill for preparing samples and for helpful conversations that facilitated the completion of this study.

