## Supplemental 1 for "Non-targeted Screening of Commercial CBD Products in the United States Reveals Common Contamination and Adulteration"

### Caffeine

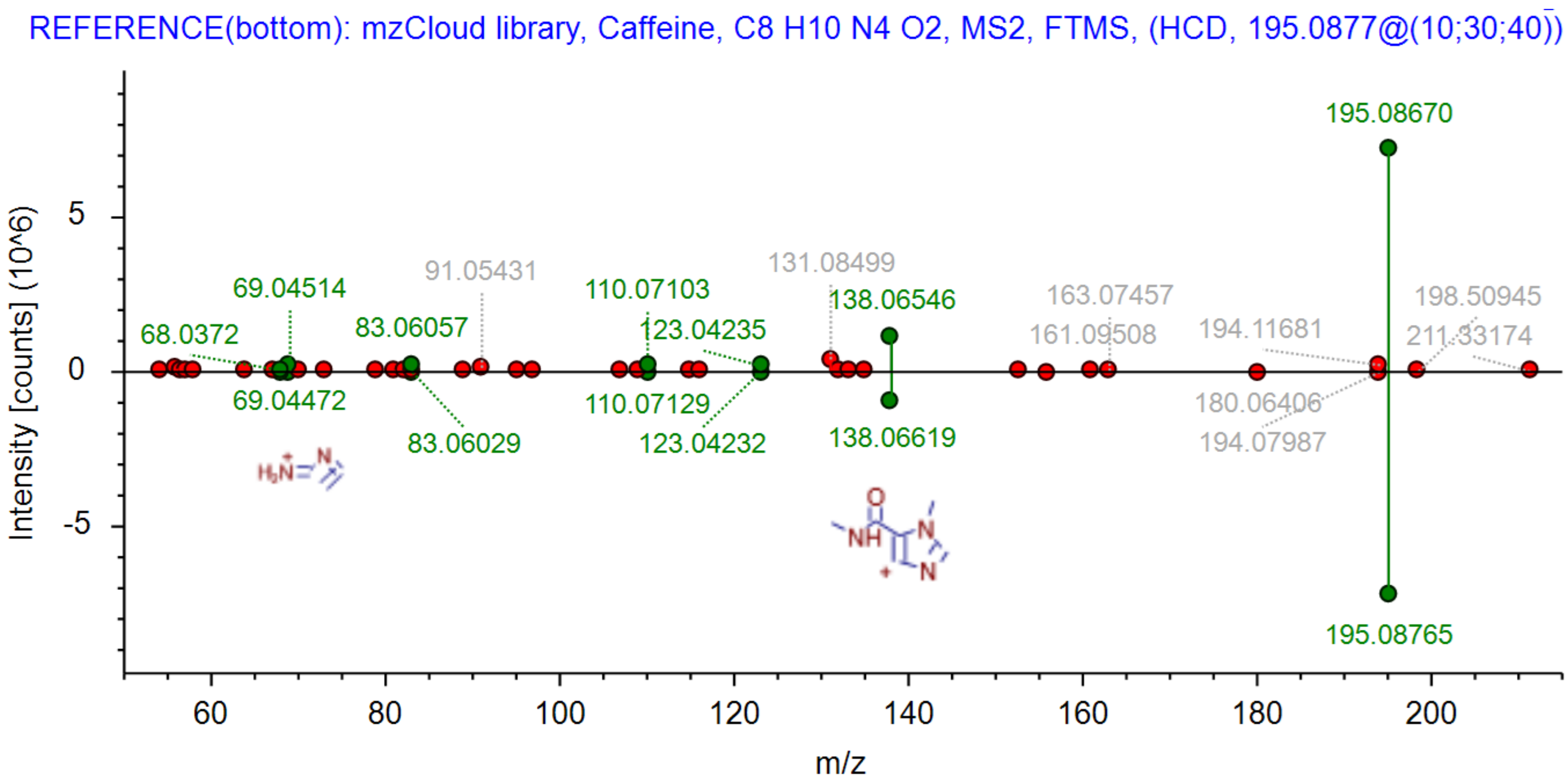

REFERENCE(bottom): mzCloud library, Caffeine, C8 H10 N4 O2, MS2, FTMS, (HCD, 195.0877@(10;30;40))
195.08670
5
0
-5
Intensity [counts] (10^6)
69.04514
68.0372
91.05431
83.06057
110.07103
131.08499
138.06546
123.04235
163.07457
161.09508
194.11681
198.50945
211.33174
69.04472
83.06029
110.07129
123.04232
138.06619
180.06406
194.07987
195.08765
60
80
100
120
140
160
180
200
m/z

### Valpromide

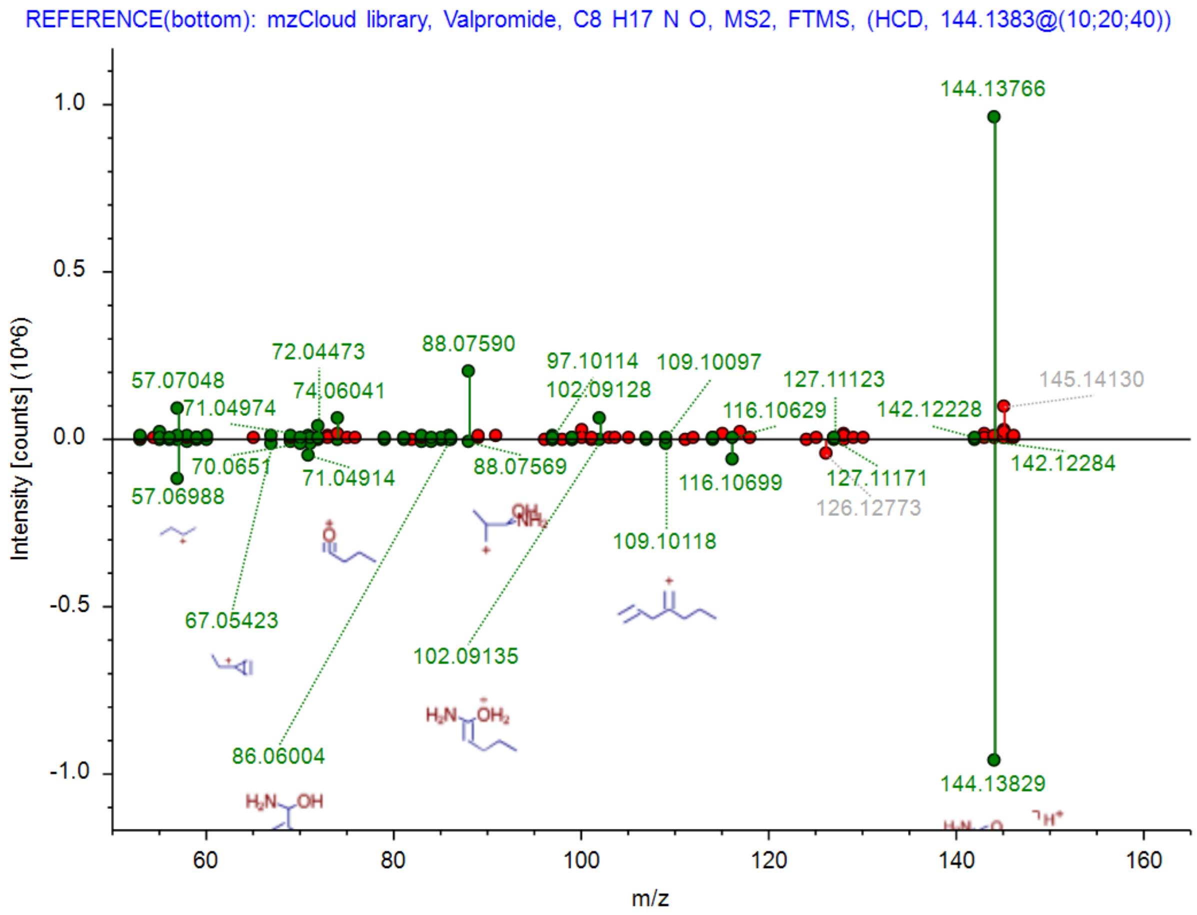

REFERENCE(bottom): mzCloud library, Valpromide, C8 H17 N O, MS2, FTMS, (HCD, 144.1383@(10;20;40))
Intensity [counts] (10^6)
1.0
0.5
0.0
-0.5
-1.0
144.13766
145.14130
57.07048
71.04974
72.04473
74.06041
88.07590
97.10114
102.09128
109.10097
116.10629
127.11123
142.12228
70.0651
71.04914
88.07569
57.06988
116.10699
127.11171
126.12773
142.12284
109.10118
67.05423
102.09135
86.06004
144.13829
60
80
100
120
140
160
m/z

### PEG N15

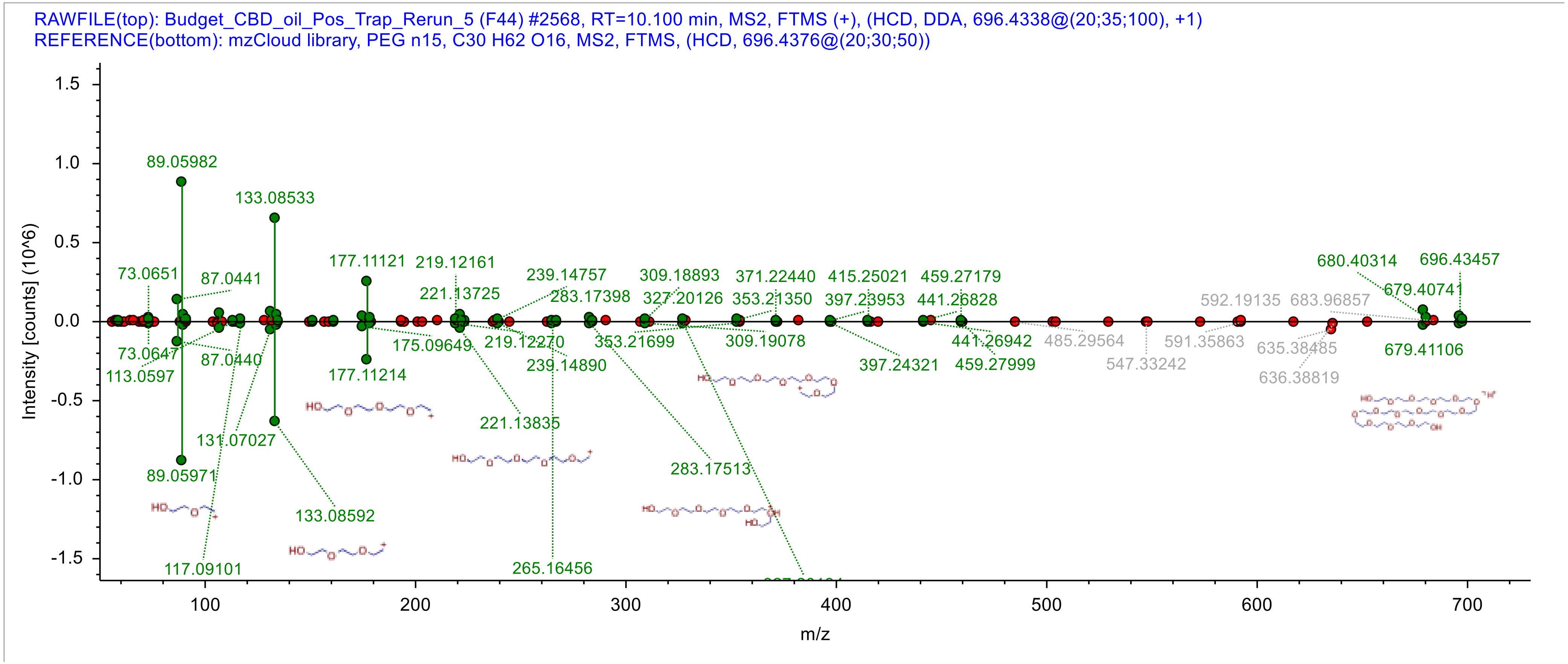
RAWFILE(top): Budget_CBD_oil_Pos_Trap_Rerun_5 (F44) #2568, RT=10.100 min, MS2, FTMS (+), (HCD, DDA, 696.4338@(20;35;100), +1)
REFERENCE(bottom): mzCloud library, PEG n15, C30 H62 O16, MS2, FTMS, (HCD, 696.4376@(20;30;50))
Intensity [counts] (10^6)
m/z
89.05982
133.08533
177.11121
219.12161
221.13725
239.14757
283.17398
309.18893
327.20126
371.22440
353.21350
415.25021
397.23953
459.27179
441.26828
680.40314
696.43457
679.40741
592.19135
683.96857
73.0651
87.0441
73.0647
87.0440
113.0597
175.09649
219.12270
239.14890
353.21699
309.19078
397.24321
441.26942
459.27999
485.29564
547.33242
591.35863
635.38485
636.38819
679.41106
177.11214
221.13835
131.07027
89.05971
133.08592
283.17513
117.09101
265.16456

### PEG N14

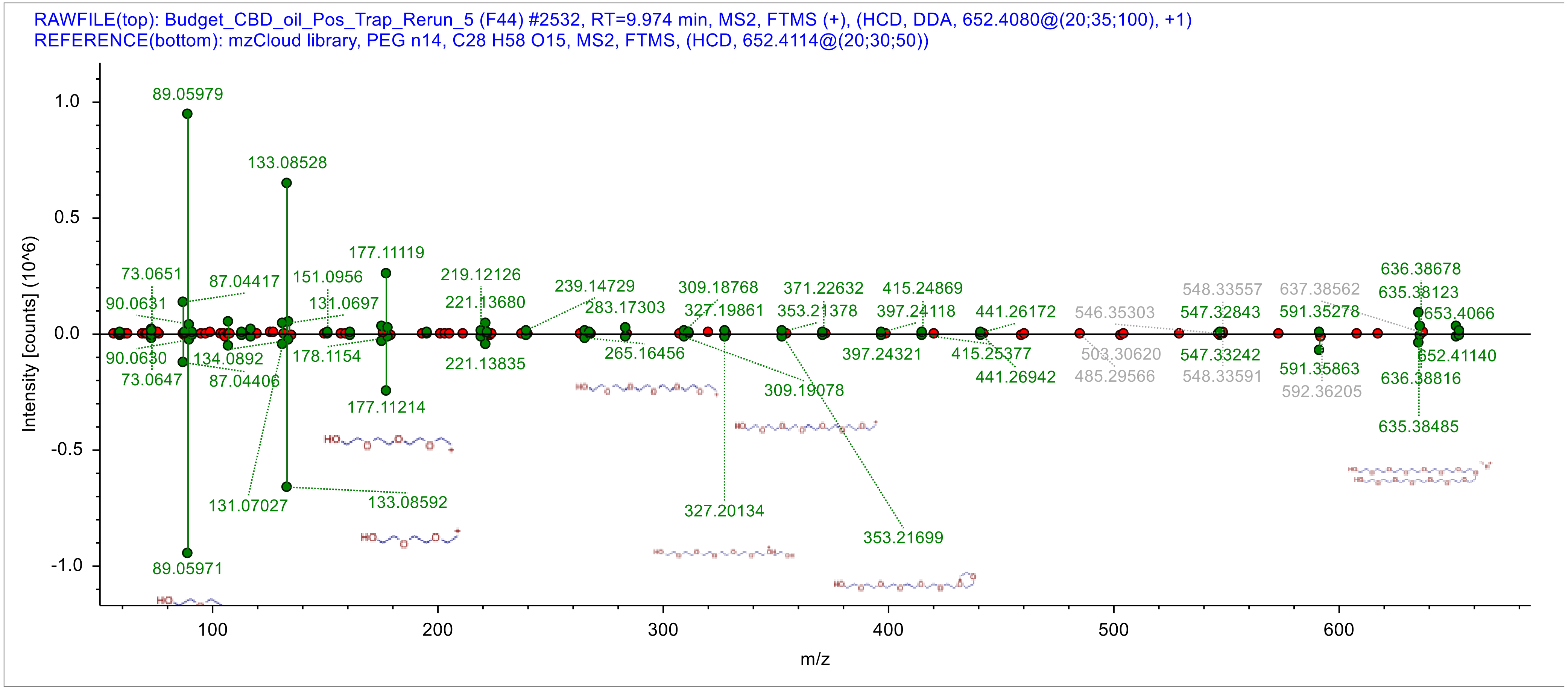
RAWFILE(top): Budget_CBD_oil_Pos_Trap_Rerun_5 (F44) #2532, RT=9.974 min, MS2, FTMS (+), (HCD, DDA, 652.4080@(20;35;100), +1)
REFERENCE(bottom): mzCloud library, PEG n14, C28 H58 O15, MS2, FTMS, (HCD, 652.4114@(20;30;50))
Intensity [counts] (10^6)
m/z
89.05979
133.08528
177.11119
73.0651
87.04417
151.0956
219.12126
239.14729
309.18768
371.22632
415.24869
636.38678
90.0631
131.0697
221.13680
283.17303
327.19861
353.21378
397.24118
441.26172
546.35303
548.33557
637.38562
635.38123
547.32843
591.35278
653.4066
90.0630
134.0892
178.1154
221.13835
265.16456
397.24321
415.25377
503.30620
547.33242
652.41140
73.0647
87.04406
441.26942
485.29566
548.33591
591.35863
636.38816
177.11214
309.19078
592.36205
635.38485
131.07027
133.08592
327.20134
353.21699
89.05971

### PEG N13

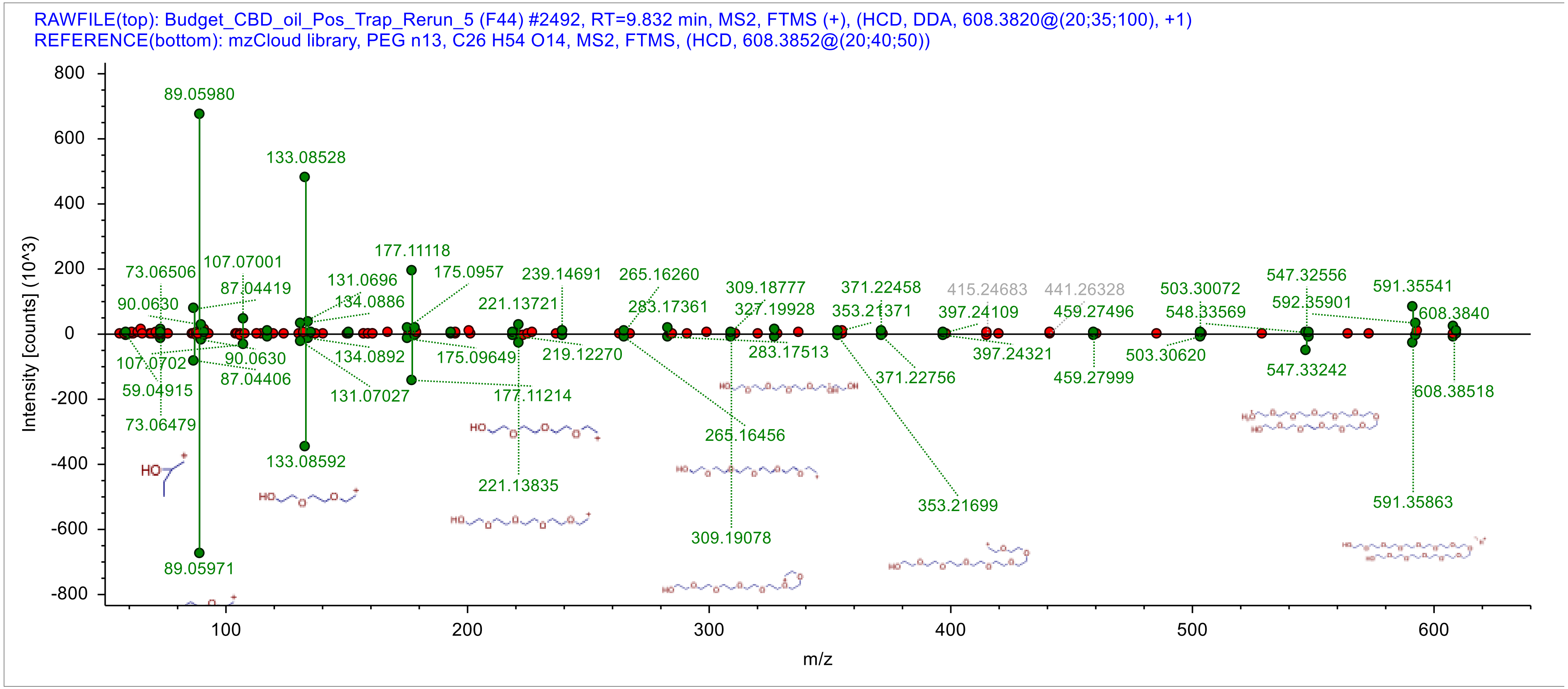

RAWFILE(top): Budget_CBD_oil_Pos_Trap_Rerun_5 (F44) #2492, RT=9.832 min, MS2, FTMS (+), (HCD, DDA, 608.3820@(20;35;100), +1)
REFERENCE(bottom): mzCloud library, PEG n13, C26 H54 O14, MS2, FTMS, (HCD, 608.3852@(20;40;50))
Intensity [counts] (10^3)
m/z
89.05980
133.08528
177.11118
175.0957
239.14691
265.16260
309.18777
371.22458
415.24683
441.26328
503.30072
547.32556
591.35541
608.3840
592.35901
548.33569
459.27496
397.24109
353.21371
327.19928
283.17361
221.13721
134.0886
131.0696
87.04419
107.07001
73.06506
90.0630
107.0702
59.04915
73.06479
90.0630
87.04406
134.0892
131.07027
175.09649
219.12270
177.11214
221.13835
283.17513
265.16456
309.19078
371.22756
353.21699
397.24321
459.27999
503.30620
547.33242
591.35863
608.38518
133.08592
89.05971

### Adipic acid

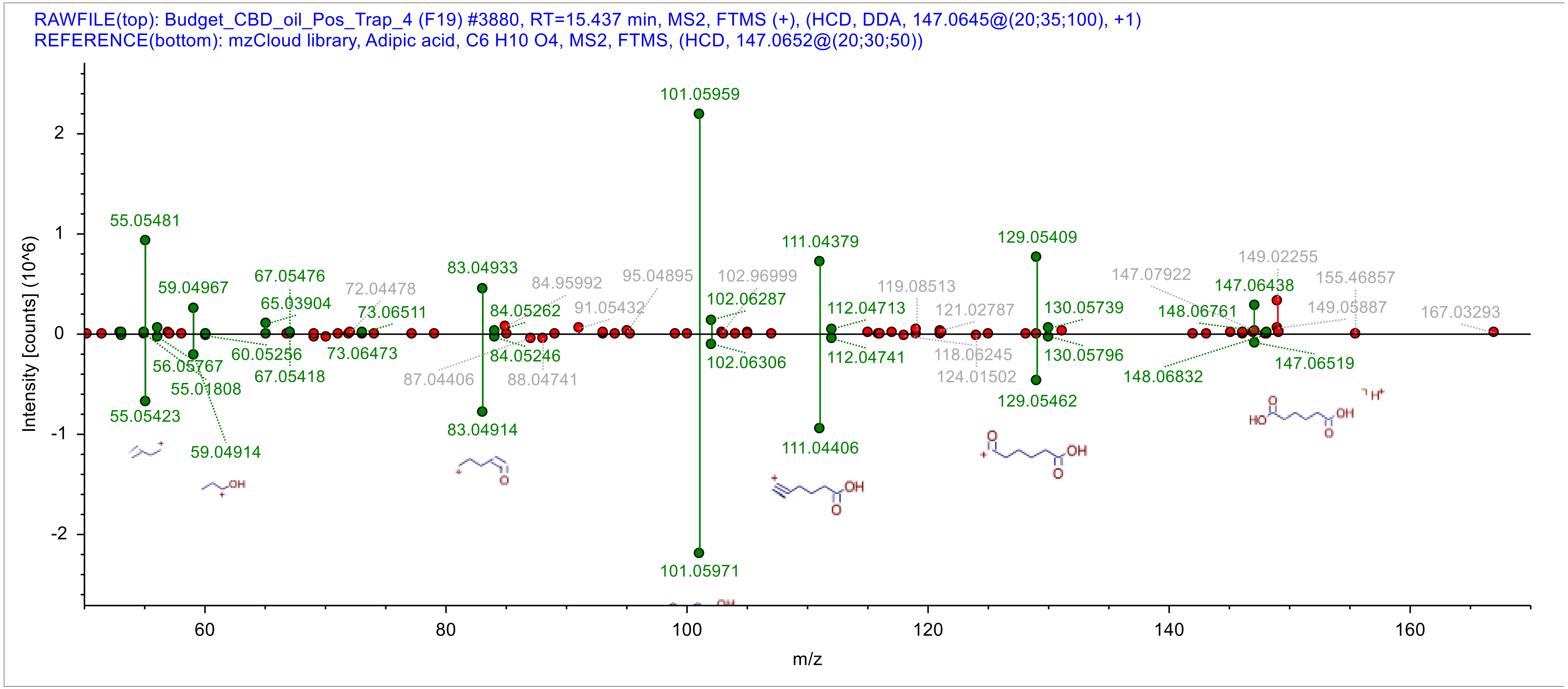

RAWFILE(top): Budget_CBD_oil_Pos_Trap_4 (F19) #3880, RT=15.437 min, MS2, FTMS (+), (HCD, DDA, 147.0645@(20;35;100), +1)
REFERENCE(bottom): mzCloud library, Adipic acid, C6 H10 O4, MS2, FTMS, (HCD, 147.0652@(20;30;50))
Intensity [counts] (10^6)
m/z
101.05959
101.05971
55.05481
55.05423
59.04967
59.04914
67.05476
65.03904
73.06511
73.06473
60.05256
56.05767
55.01808
67.05418
83.04933
83.04914
84.05262
84.05246
84.95992
91.05432
95.04895
87.04406
88.04741
102.96999
102.06287
102.06306
111.04379
111.04406
112.04713
112.04741
119.08513
121.02787
118.06245
124.01502
129.05409
129.05462
130.05739
130.05796
147.07922
148.06761
148.06832
147.06438
147.06519
149.02255
155.46857
149.05887
167.03293

### Palmitoleic acid

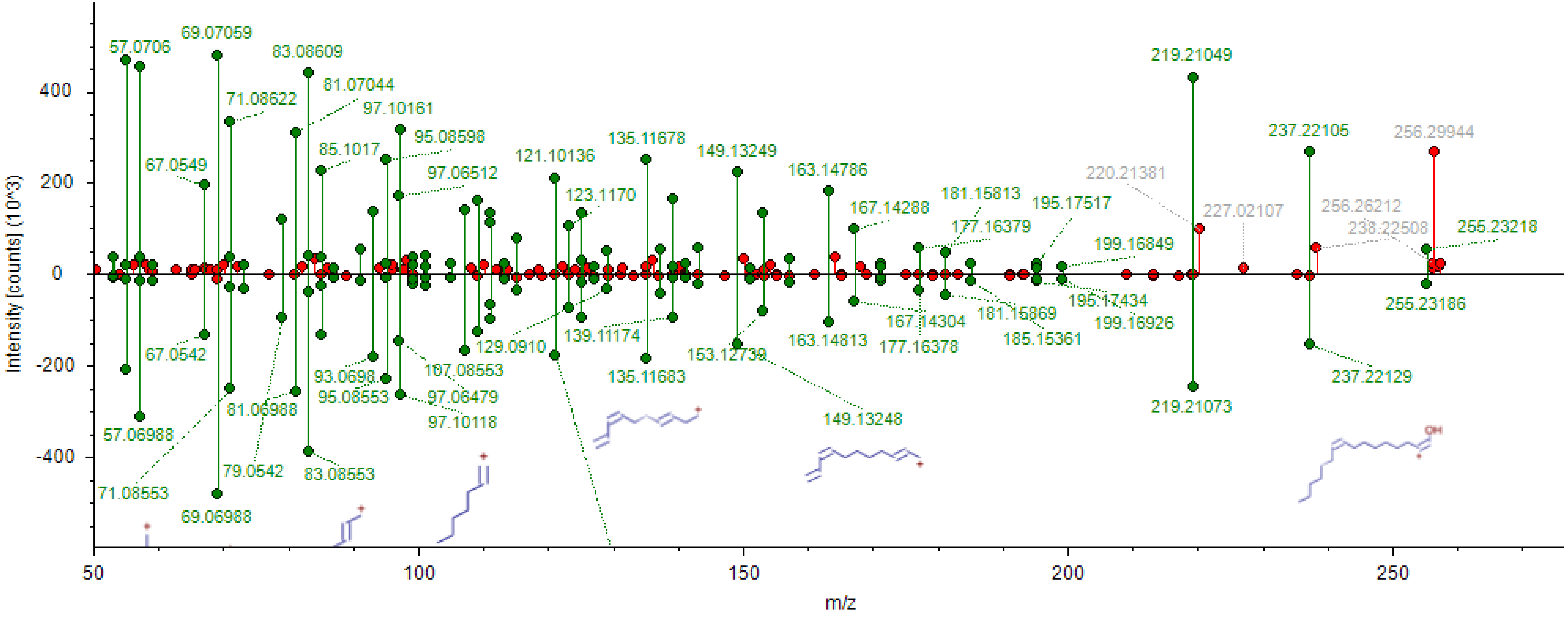

### Melatonin

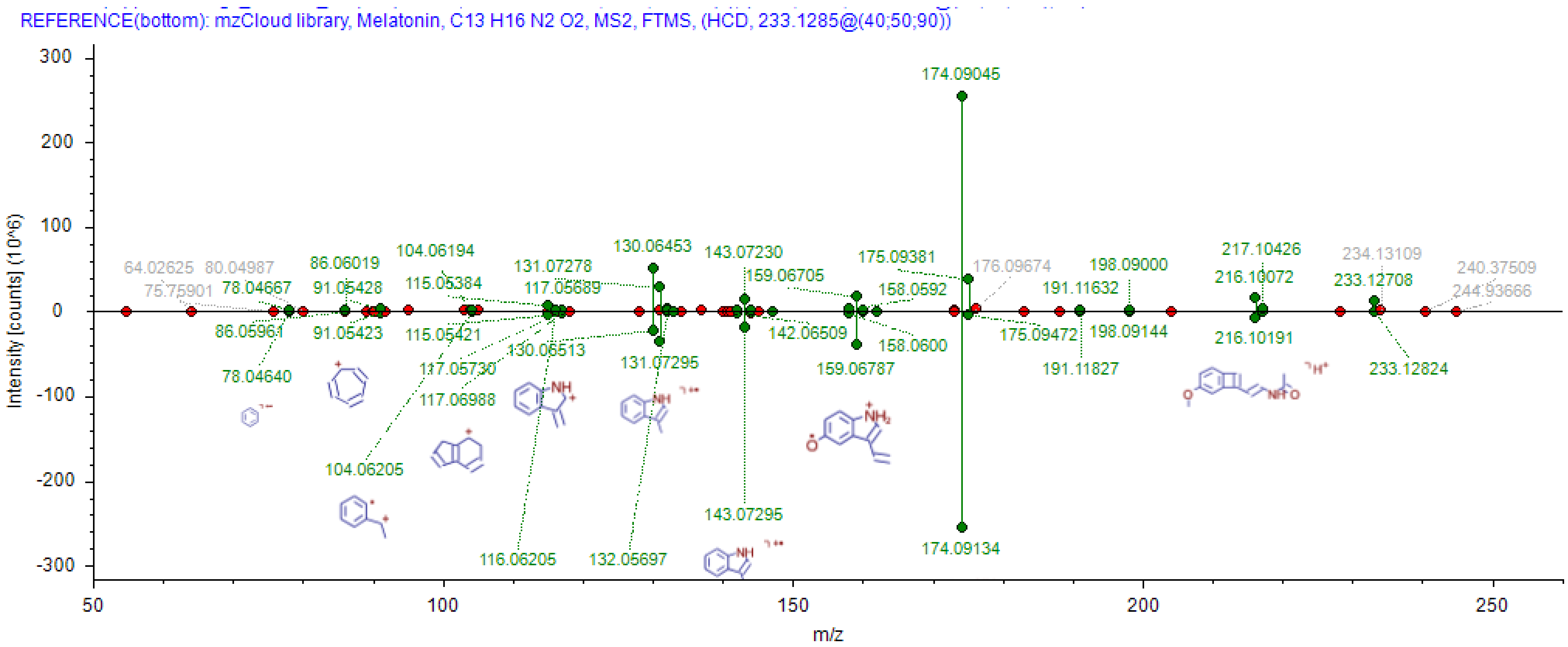
REFERENCE(bottom): mzCloud library, Melatonin, C13 H16 N2 O2, MS2, FTMS, (HCD, 233.1285@(40;50;90))
Intensity [counts] (10^6)
300
200
100
0
-100
-200
-300
m/z
50
100
150
200
250
174.09045
64.02625
80.04987
75.75901
78.04667
86.06019
91.05428
104.06194
115.05384
131.07278
117.05689
130.06453
143.07230
159.06705
175.09381
158.0592
176.09674
198.09000
191.11632
217.10426
216.10072
234.13109
233.12708
240.37509
244.93666
86.05961
91.05423
115.05421
130.06513
131.07295
142.06509
158.0600
175.09472
198.09144
216.10191
78.04640
117.05730
117.06988
159.06787
191.11827
233.12824
104.06205
116.06205
132.05697
143.07295
174.09134

### Theobromine

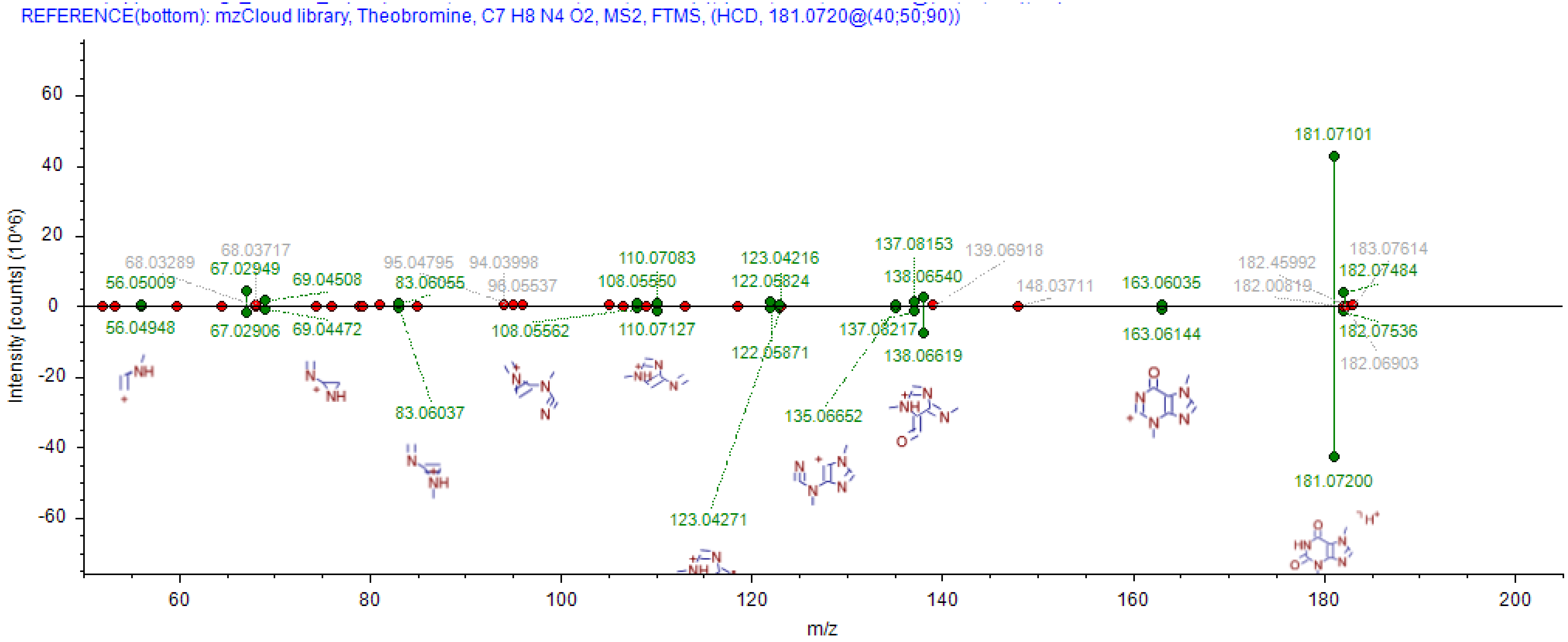

### 1-Napthol

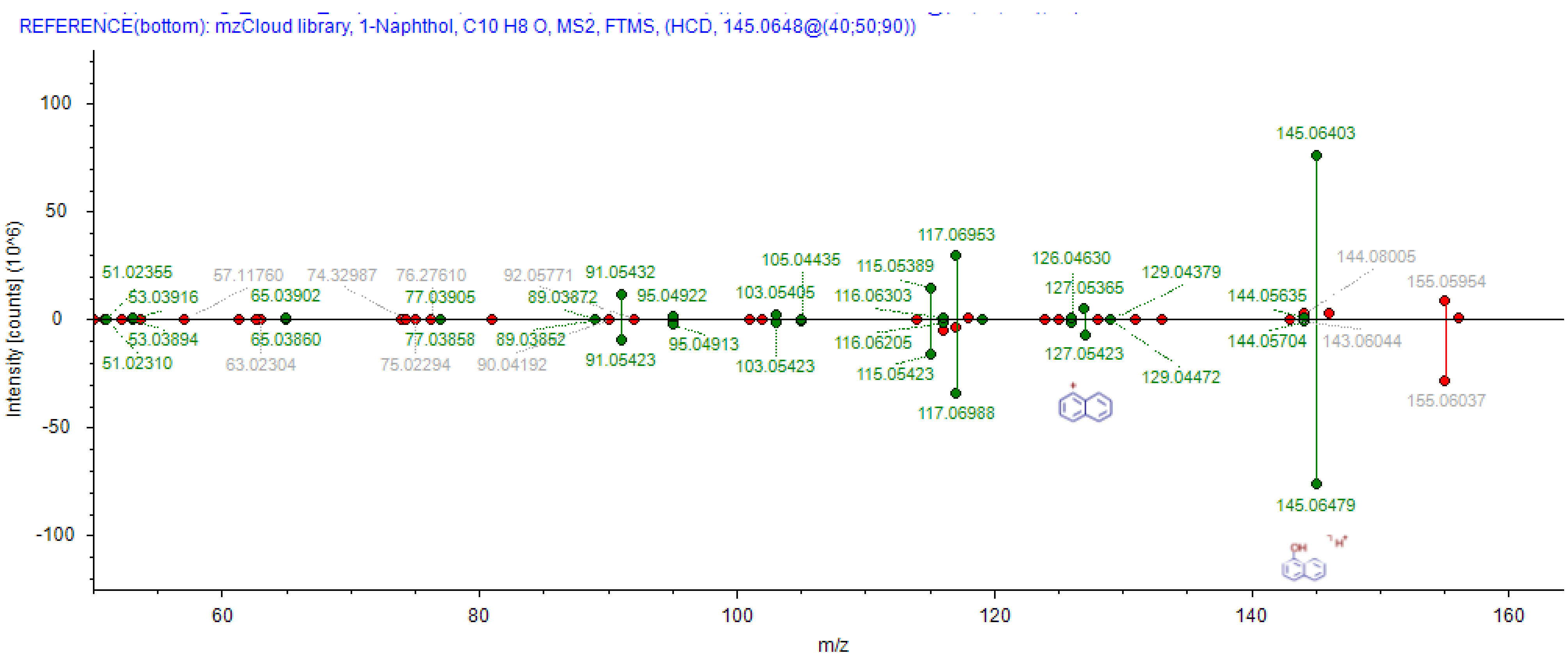

REFERENCE(bottom): mzCloud library, 1-Naphthol, C10 H8 O, MS2, FTMS, (HCD, 145.0648@(40;50;90))
Intensity [counts] (10^6)
m/z
100
50
0
-50
-100
60
80
100
120
140
160
145.06403
145.06479
117.06953
117.06988
105.04435
103.05405
103.05423
115.05389
115.05423
116.06303
116.06205
126.04630
127.05365
127.05423
129.04379
129.04472
144.05635
144.05704
144.08005
143.06044
155.05954
155.06037
51.02355
51.02310
53.03916
53.03894
57.11760
65.03902
65.03860
63.02304
74.32987
76.27610
77.03905
77.03858
75.02294
92.05771
89.03872
89.03852
90.04192
91.05432
91.05423
95.04922
95.04913

### Nootkatone

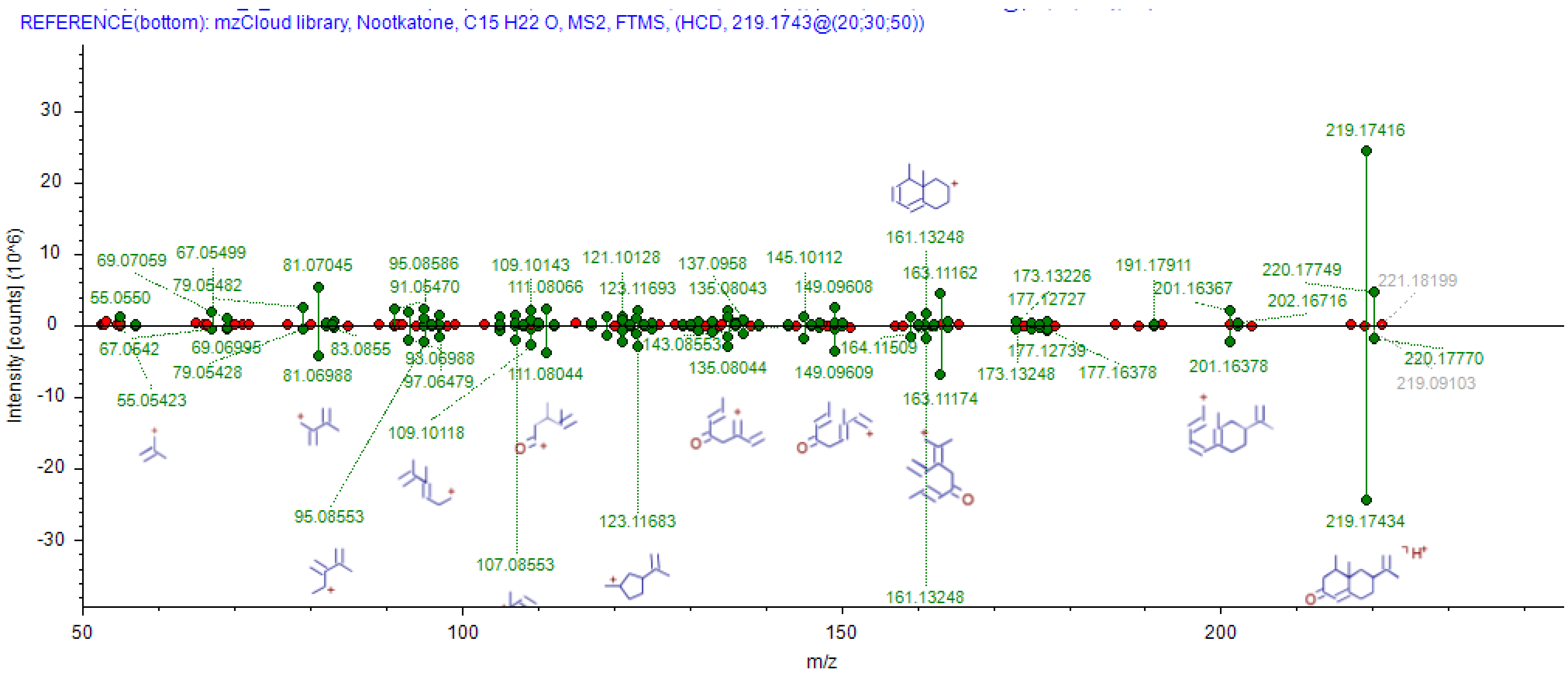
REFERENCE(bottom): mzCloud library, Nootkatone, C15 H22 O, MS2, FTMS, (HCD, 219.1743@(20;30;50))
Intensity [counts] (10^6)
m/z
219.17416
219.17434
161.13248
163.11162
163.11174

### Yohimbine

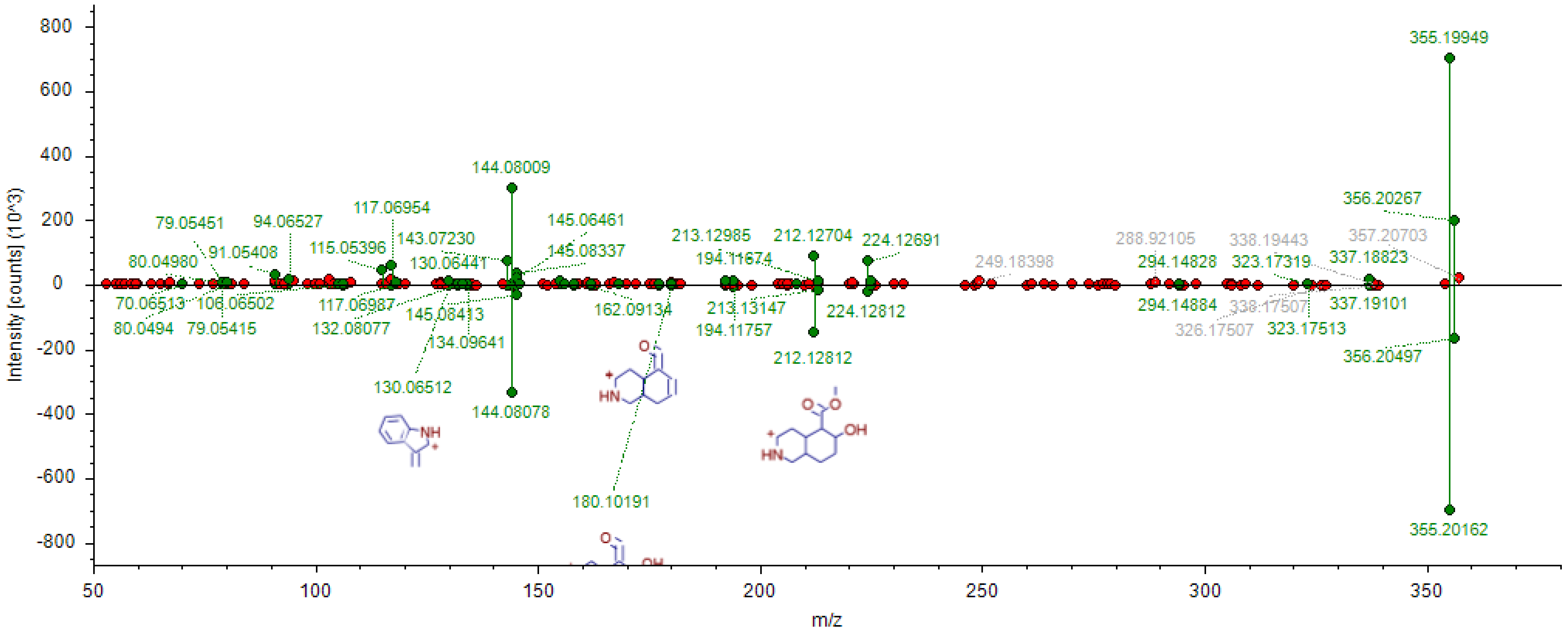

355.19949
144.08009
356.20267
117.06954
145.06461
79.05451
94.06527
143.07230
213.12985
212.12704
224.12691
288.92105
338.19443
357.20703
91.05408
115.05396
145.08337
194.11674
249.18398
294.14828
323.17319
337.18823
80.04980
130.06441
70.06513
106.06502
117.06987
162.09134
213.13147
224.12812
294.14884
338.17507
337.19101
80.0494
79.05415
132.08077
145.08413
194.11757
326.17507
323.17513
134.09641
212.12812
356.20497
130.06512
144.08078
180.10191
355.20162
Intensity [counts] (10^3)
m/z

### Erucamide

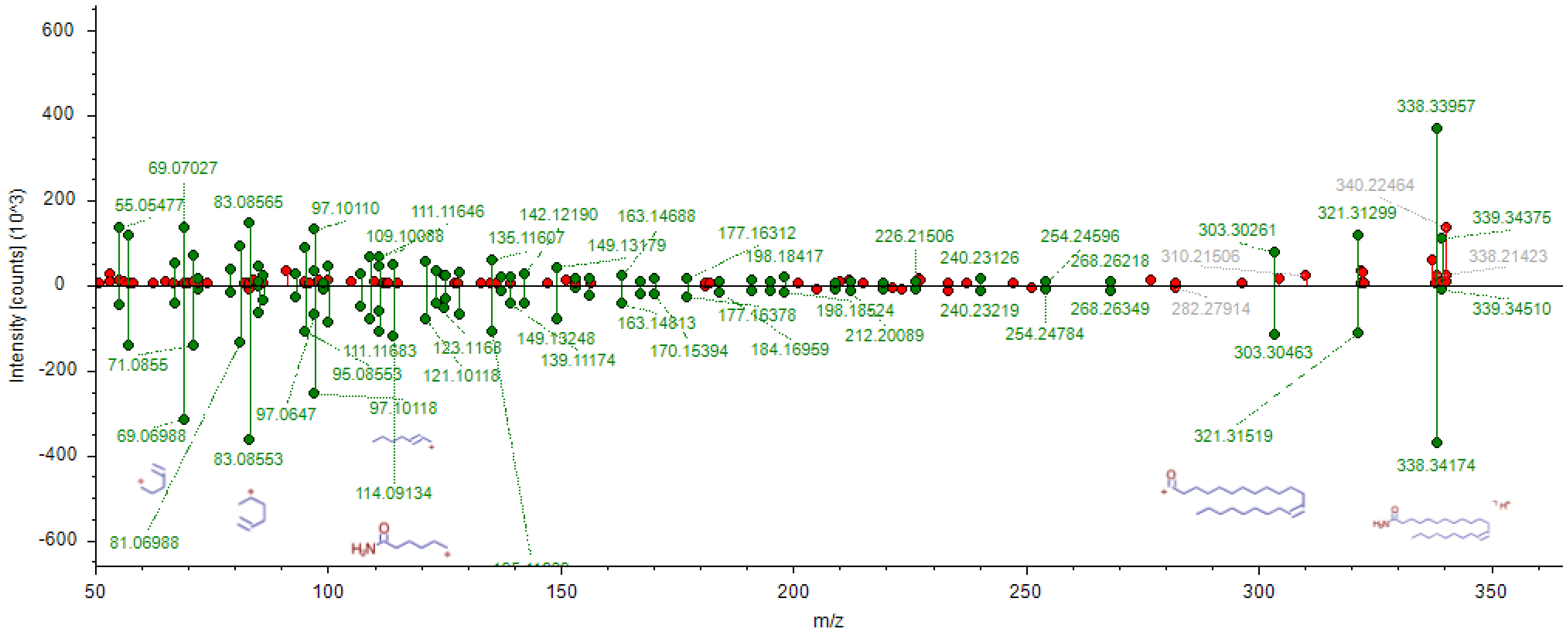

600
400
200
0
-200
-400
-600
Intensity [counts] (10^3)
50
100
150
200
250
300
350
m/z
338.33957
69.07027
55.05477
83.08565
97.10110
111.11646
109.10088
142.12190
135.11607
163.14688
149.13179
177.16312
198.18417
226.21506
240.23126
254.24596
268.26218
310.21506
303.30261
321.31299
340.22464
339.34375
338.21423
339.34510
71.0855
69.06988
83.08553
81.06988
97.0647
95.08553
111.11683
97.10118
114.09134
123.1168
121.10118
149.13248
139.11174
163.14813
170.15394
177.16378
184.16959
198.18524
212.20089
240.23219
254.24784
268.26349
282.27914
303.30463
321.31519
338.34174
